# Exploiting the Affimer platform against influenza A virus

**DOI:** 10.1101/2023.08.22.554342

**Authors:** Oliver Debski-Antoniak, Alex Flynn, David P. Klebl, Christian Tiede, Ian A. Wilson, Stephen P. Muench, Darren Tomlinson, Juan Fontana

**Affiliations:** School of Molecular and Cellular Biology, Faculty of Biological Sciences, University of Leeds, LS2 9JT, United Kingdom; Astbury Centre for Structural and Molecular Biology, University of Leeds, LS2 9JT, United Kingdom; School of Biomedical Sciences, Faculty of Biological Sciences, University of Leeds, LS2 9JT, United Kingdom; Department of Integrative Structural and Computational Biology, The Scripps Research Institute, La Jolla, CA 92037, USA

**Keywords:** influenza A virus, Affimer, mAb, cryo-electron microscopy sample preparation

## Abstract

Influenza A virus (IAV) is well known for its pandemic potential. While current surveillance and vaccination strategies are highly effective, therapeutic approaches are short-lived due to the high mutation rates of IAV. Currently, monoclonal antibodies (mAbs) have emerged as a promising approach to tackle future IAV pandemics. Additionally, several antibody-like alternatives exist that aim to improve upon mAbs. Affimers, one such alternative, benefit from a short development time, high expression levels in *E. coli*, and complete animal-free production. Here we exploited the Affimer platform to isolate and produce specific and potent inhibitors of IAV. Starting from a monomeric version of the IAV trimeric hemagglutinin (HA) fusion protein, we isolated 12 Affimers that inhibit IAV H3 subtype infection *in vitro*. Two of these Affimers were characterised in detail: they exhibited binding affinities to the target H3 HA protein in the nM range and bound specifically to the HA1 head domain. Cryo-EM employing a novel spray approach to prepare cryo-grids allowed us to image HA-Affimer complexes. Combined with functional assays, we determined that the mode of inhibition of these Affimers is based on blocking the interaction of HA to the host-cell receptor - sialic acid. Additionally, these Affimers inhibited IAV strains closely related to the one employed for Affimer isolation. Overall, these results support the use of Affimers as an alternative to existing targeted therapies for IAV and pave the way for their use as diagnostic reagents.

**Importance:** Influenza A virus is one of the few viruses that can cause devastating pandemics. Due to the high mutation rates of this virus, annual vaccination is required and antivirals are short-lived. Monoclonal antibodies present a promising approach to tackle influenza virus infections, but are associated with some limitations. To improve on this strategy, we explored the Affimer platform, which are antibody-like, bacterially made proteins. By performing phage-display against a monomeric version of influenza virus fusion protein, an established viral target, we were able to isolate Affimers that inhibit influenza virus infection *in vitro*. We characterised the mechanism of inhibition of the Affimers by challenging with related influenza virus strains. We additionally characterised an HA-Affimer complex structure, using a novel approach to prepare samples for cryo-electron microscopy. Overall, these results show that Affimers are a promising tool against influenza virus infection.

## Introduction

Influenza A virus (IAV) is a paradigmatic virus with pandemic potential. The ability of two different IAV strains to recombine and reassort their viral RNA segments/RNPs and generate a novel strain (a process termed antigenic shift), enables IAV to spontaneously undergo dramatic changes in the nature and composition of key proteins in the virus, which combined with its large host range can result in worldwide zoonotic infection (1). Additionally, the error-prone nature of IAV RNA-dependant RNA polymerase enables the virus to further evolve antigenically (antigenic drift) (2), ultimately resulting in acquired resistance to licensed antivirals and pre-existing immunity, and the necessity for annual vaccination programmes to maintain immunity in the vulnerable to prevent serious disease (3). Previous IAV pandemics include the 1918 avian pandemic (the Spanish flu), which is estimated to have claimed more than 50 million lives and caused more than 500 million infections worldwide (4), and more recently, the 2009 swine flu pandemic (5). Given the antigenic shift and drift of IAV, it is likely that further IAV pandemics will occur in the future. For example, both H5N1 and H7N9 of avian origins have been reported to have jumped into humans in multiple instances, with mortalities reported to be as high as 60% and 40%, respectively (6,7). Due to the nature of these zoonotic outbreaks alongside the detrimental consequences, improved surveillance to rapidly identify and isolate these spill-over events, combined with a coordinated therapeutic effort, will be critical to prevent the impact that future IAV pandemics will have on society.

Neutralizing monoclonal antibodies (mAbs) used as diagnostic and therapeutic approaches are one of the most promising tools for tackling IAV pandemics, in conjunction with annual vaccination campaigns (8). However, mAbs are expensive and often difficult to develop and manufacture, meaning low and middle income countries cannot access or afford such therapies (9), thus preventing an effective solution to a global problem. This highlights the need for alternative tools and strategies to overcome the practical and logistical limitations of mAbs that can be utilised alongside current treatments to add to the arsenal of approaches to mitigate against future pandemiccs.

Recently, small, synthetic antibody-like proteins termed Affimers have become available as emerging reagents for diagnostics and biotherapeutics (10–12). For example, Affimers have been employed in diagnostic tests for detecting the human-infecting Crimean-Congo hemorrhagic fever virus and the plant-infecting Cowpea Mosaic virus (13,14), and are actively being developed as oncological therapeutics. Affimer molecules are small (∼12 kDa) proteins based on naturally occurring cystatins (cysteine protease inhibitors). These molecules display stability up to high temperatures (∼ 100 °C) (12), removing the need for cold chain distribution around the globe. Affimer reagents display two variable peptide regions that allow binding to a target in a similar manner to mAbs, with affinities typically in the low nM range (15). Affimer isolation is performed via phage-display, followed by expression in bacteria. This not only prevents any ethical concerns by removing the need for animals, but also facilitates rapid, cost-effective, and reproducible production. Here we report the first isolation and characterisation of Affimer molecules against IAV.

Hemagglutinin (HA), one of two glycoproteins of IAV, is responsible for virus entry into the host cell. HA comprises two domains: 1) HA1, which includes the N-terminal domain (NTD) and the receptor binding domain (RBD), is responsible for interaction with host receptors such as sialic acid and entry into host cells; 2) HA2, which along with N- and C-termini of HA1 constitutes the fusion domain, is responsible for virus-host cell membrane fusion through conformational changes triggered by the low pH within endocytic compartments that IAV hijacks. In addition to interactions with sialic acid and membrane fusion, HA is also the most abundant glycoprotein in infectious influenza virions and is also involved in virus assembly and egress. Due to its multifunctionality, HA is an attractive drug target, as its different functions can be inhibited independently, presenting several target sites on a single viral protein. As such, HA has become a validated therapeutic target (16). Whilst HA is extremely desirable as a therapeutic target, it poses many challenges. For example, HA has evolved to support a high degree of mutations resulting from immunogenic pressure. The breadth of host-species has enabled further inter-subtype variability and the spike is heavily decorated with glycans, protecting vulnerable epitopes from antibodies and bulky therapeutics. To date, there is only one known mAb (CR9114) with the capacity to universally recognise group 1 and 2 HA proteins alongside influenza B virus HA spikes (17), although this mAb is incapable of neutralising all viruses from different strains and subtypes due to the high antigenic variability in the HAs.

Here we describe the isolation of Affimer molecules specific to the IAV spike protein HA. Two of these Affimers were characterised and found to be potent IAV inhibitors through their binding to the HA RBD and blocking its interaction with sialic acid. This mechanism of inhibition was further supported by cryo-electron microscopy (cryo-EM) of HA-Affimer complexes. Importantly, we demonstrate that Affimer molecules can inhibit virus efficiently and show their breadth in tackling variants generated through antigenic drift.

## Results

### Isolation of Affimer molecules against IAV HA

To assess the potential for Affimers as tool compounds against IAV, Affimer molecules were isolated via phage display, expressed in *E. coli*, and characterised (Fig. 1 and Sup. Fig. 1). Briefly, we immobilised a monomer of the trimeric spike protein HA from the pandemic A/Aichi/68 (H3N2) virus and a subset of 480 phage clones were selected after three phage-display panning rounds. The individual phages were prepared for initial evaluation. Target binding was carried out via phage enzyme-linked immunosorbent assay (ELISA) (12), and a threshold of > 0.5 (absorbance at 620 nm) was employed for selection, resulting in 192 candidates (Sup. Fig. 1A). The DNA sequences of the Affimer binders were determined, resulting in 34 unique Affimer sequences (Sup. Figs. 1B and 2). None of these Affimers were cytotoxic for tissue cultured cells when incubated at 100 µM for 72 hrs (Sup. Fig. 1C). A microneutralisation assay determined that seven of these Affimers inhibited IAV infection *in vitro*, when added to tissue cultured cells at 3.57 µM (50 µg/mL; Sup. Fig. 1D). Herein we describe the characterisation of two of the promising candidates from this pool: A5 and A31.

**Figure 1.**
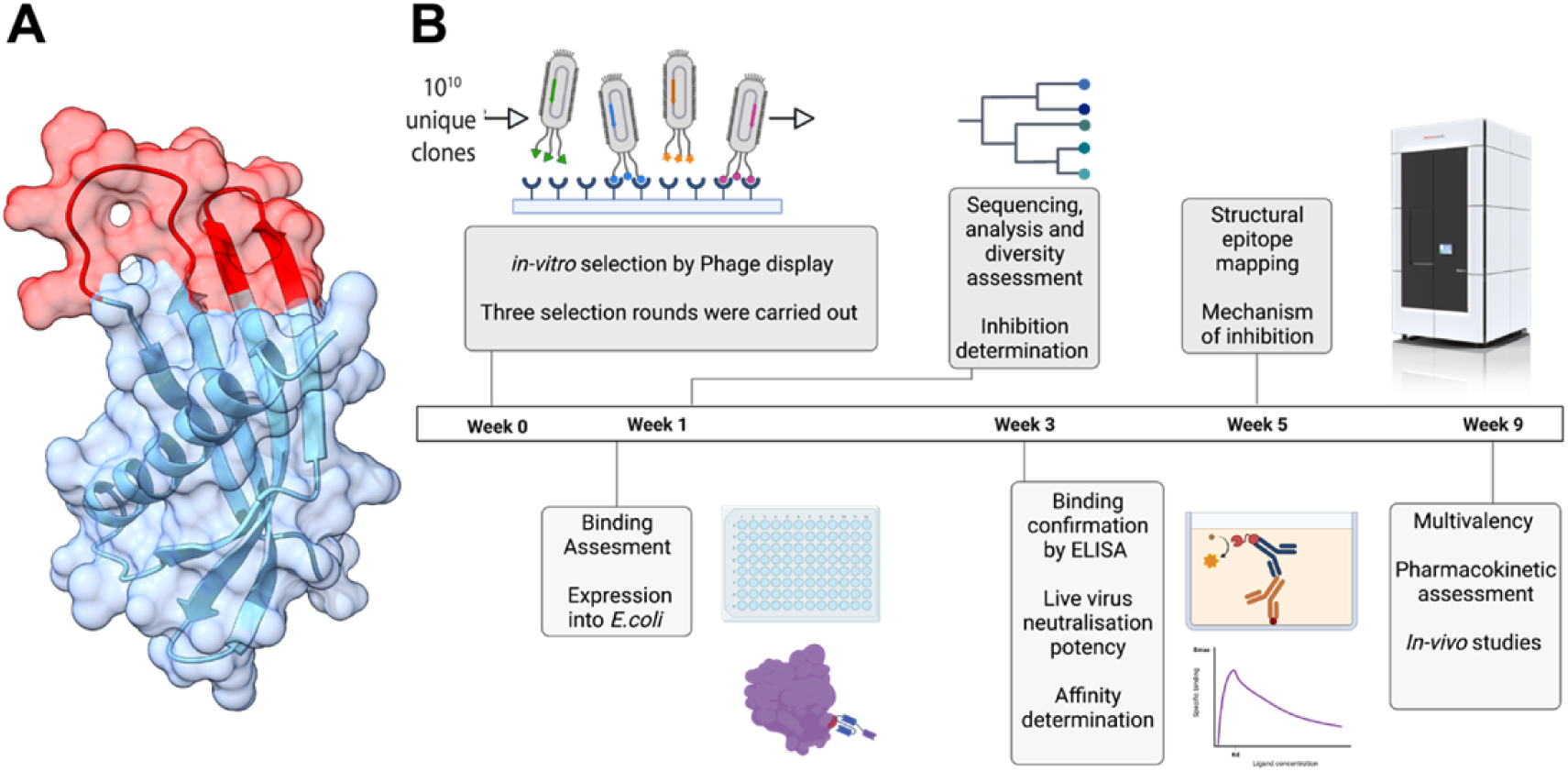
Workflow for isolating and characterising Affimer molecules against IAV HA. **(A)** Atomic model of an Affimer molecule, overlapped with its surface representation. The two variable peptide regions are highlighted in red. **(B)** Workflow overview for the generation of anti-IAV Affimer molecules. Top-left to bottom-right, schematics represent a phage-ELISA, phylogenetic tree, structure determination by cryo-electron microscopy, binding assessment by ELISA, and affinity assessment via ELISA and SPR.

### Affimer molecules show high affinity against target protein HA

Affimers A5 and A31 were selected due to their high level of neutralisation (Sup. Fig. 1D) and their high sequence diversity within the neutralising Affimers, as determined by a phylogenetic tree of the variable region of isolated Affimers (Sup. Fig. 2). To determine their binding affinities, surface plasmon resonance (SPR) spectroscopy and ELISAs were employed, again utilising a monomeric HA [A/Aichi/68 (H3N2)] (Fig. 2). For SPR, Affimer molecules were immobilised onto a streptavidin-coated chip before a range of concentrations of monomeric HA were flowed over the Affimers. A5 and A31 showed Kd values in the low nM range (2.80 ± 1.15 nM for A5 and 5.94 ± 3.27 nM for A31; Figs. 2A and B), demonstrating high affinity interactions. Both A5 and A31 displayed relatively slow K_on_ association rates (1.15 x 10^5^ M^-1^ s^-1^ for A5 and 2.15 x 10^4^ M^-1^ s^-1^ for A31) but very slow K_off_ rates of dissociation (1.72 x 10^-4^ s^-1^ for A5 and 1.92 x 10^-4^ s^-1^ for A31), indicating slow but tight binding of these molecules. Assessment by ELISA further confirmed high affinity binding to HA. Mean IC_50_ values were calculated by ELISA as 1.03 nM for A5 and 1.10 nM for A31 (Fig. 2C), in the range of the Kd values determined by SPR. In conclusion, both A5 and A31 bind to their target protein HA with high affinities, comparable to those for mAbs raised against HA (0.1 nM-500 nM)(18–21).

**Figure 2.**
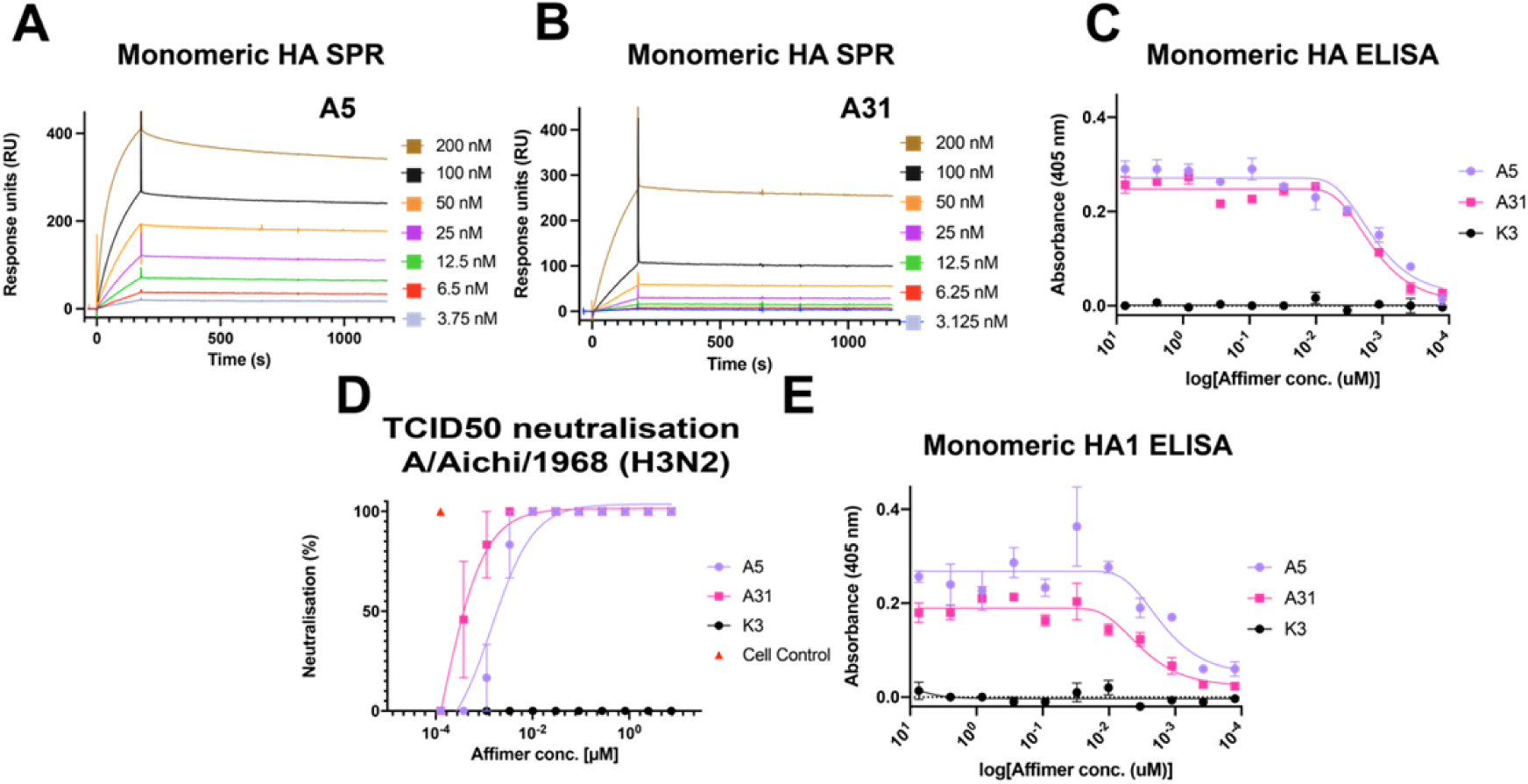
Affimer affinity for IAV monomeric HA and Affimer neutralisation efficiency (A-B) Binding of monomeric HA to Affimers was assessed via SPR. HA, in a concentration range (3.75 nM-200 nM), was flown over either immobilised A5 **(A)** or A31 **(B)**. As monomeric HA was utilised, the HA sensograms were fitted to a Langmuir model, assuming a 1:1 binding interaction. **(C)** An ELISA was employed as an alternative method to determine the binding affinities of the Affimer molecules to HA. Monomeric *[A/Aichi/1968 (H3N2)]* HA was absorbed to Maxisorb plates, before a concentration range (7.41 μM-125.5 pM) of biotinylated anti-HA Affimer (A5 (purple circles), A31(pink squares) or non-specific K3 (black circles); the same colour scheme follows throughout). All conditions were carried out in triplicate. ELISA binding curves from experimental triplicates are shown with one site—specific binding curves fit to the data. **(D)** 100 x TCID_50_ assay in which A/Aichi/1968 (H3N2) was incubated with either A5 or A31 in a concentration-dependant manner (7.41 μM-125.5 pM). **(E)** ELISA to assess binding of A5 or A31 to immobilised HA1 from A/Aichi/1968 (H3N2) (7.41 μM-125.5 pM). All assays were completed with two biological repeats. In C-E, data are mean and error bars represent standard deviation (from two biological repeats, each performed in triplicate).

### Affimer molecules show strong potency against IAV through interaction with the receptor binding site

Once we confirmed that Affimers A5 and A31 bind to HA, we explored their neutralisation potency through a TCID_50_ assay against IAV *[A/Aichi/1968 (H3N2)].* In this assay, cultured cells were infected with IAV at a concentration that would result in death of the whole monolayer. Affimers were then added to the virus prior to infection in a concentration-dependent manner to monitor inhibition. If Affimers inhibit IAV infection, cells would remain alive, and an intact monolayer would be observed. Both A5 and A31 showed comparable, high potency against IAV in the low nM range. The resulting TCID_50_ values were 1.96 nM and 0.44 nM, respectively (Fig. 2D and Sup. Fig. 3A), showing comparable potencies to those of mAbs (22–24).

To map the epitopes to which both Affimers bind, an ELISA was employed again. In this instance, monomeric HA1, the receptor-binding subunit of HA, was immobilised to wells of a MaxiSorp plate before being incubated with either A5, A31 or control, in a concentration-dependent manner in an ELISA (Fig. 2E). Mean IC_50_ values of 0.93 nM and 2.45 nM respectively for A5 and A31 were obtained (Fig. 2E), which are comparable to those of full-length HA (Fig. 2C), suggesting that the binding of these Affimer molecules is strictly localised to the HA1 head domain.

To further identify the HA epitopes of A5 and A31, cryo-EM was carried out on each HA- Affimer complex using a trimeric form of HA (HA-HK/68 H3), which is 98.6% sequence identical to the HA from *A/Aichi/1968* used to isolate the Affimers (25). To overcome the preferred orientation of HA observed when grids were prepared using the traditional plunge-freezing vitrification approach (26), we adopted rapid-spray and vitrification for grid preparation (Sup. Fig. 4 and Sup. Table. 1) (26,27), which enabled an H3 HA-only cryo-EM average at 4.3 Å resolution (Sup. Figs. 4A & B and 5). In addition to a preferred orientation, we noticed that A5 induced HA aggregation when HA-A5 grids were prepared by plunge-freezing. To overcome the on-grid HA aggregation induced by A5, we combined the spray-based grid preparation (27) with rapid mixing (Sup. Figs. 4C and D), with the aim of trapping HA after Affimer binding but before aggregation. On the other hand, HA and A31 were pre-incubated for 15 minutes at room temperature, prior to spraying (Sup. Figs. 4E and F). This led to reconstructions of HA in complex with A5 and A31 to global resolutions of 4.4 Å and 3.4 Å, respectively, with imposed C3 symmetry (Figs. 3A and B). The resulting cryo-EM averages confirmed biochemical observations and revealed both A5 and A31 interact either directly with or close to the RBD of HA, potentially obstructing cell receptor entry (Fig. 3A and B). The resolution for the H3-A5 complex was not high enough for accurate model building, likely due to a combination of A5-induced aggregation, preferred orientation, and limited particle numbers, ultimately leading to strong anisotropy in the map, despite our best efforts to reduce this phenomenon. Rapid mixing reduced aggregation but did not prevent it, reducing the number of particles in a range of orientations available for final reconstructions (23,369 particles vs 104,544 particles in H3-A31) (Sup. Figs. 6 and 7). Therefore, an H3 trimer (PDB ID: 4FNK) and three A5 homology models were rigid-body fitted into a deepEMhancer-filtered H3-A5 map. However, the H3-A31 complex was resolved to sufficient resolution to enable model building into the EM density.

**Figure 3.**
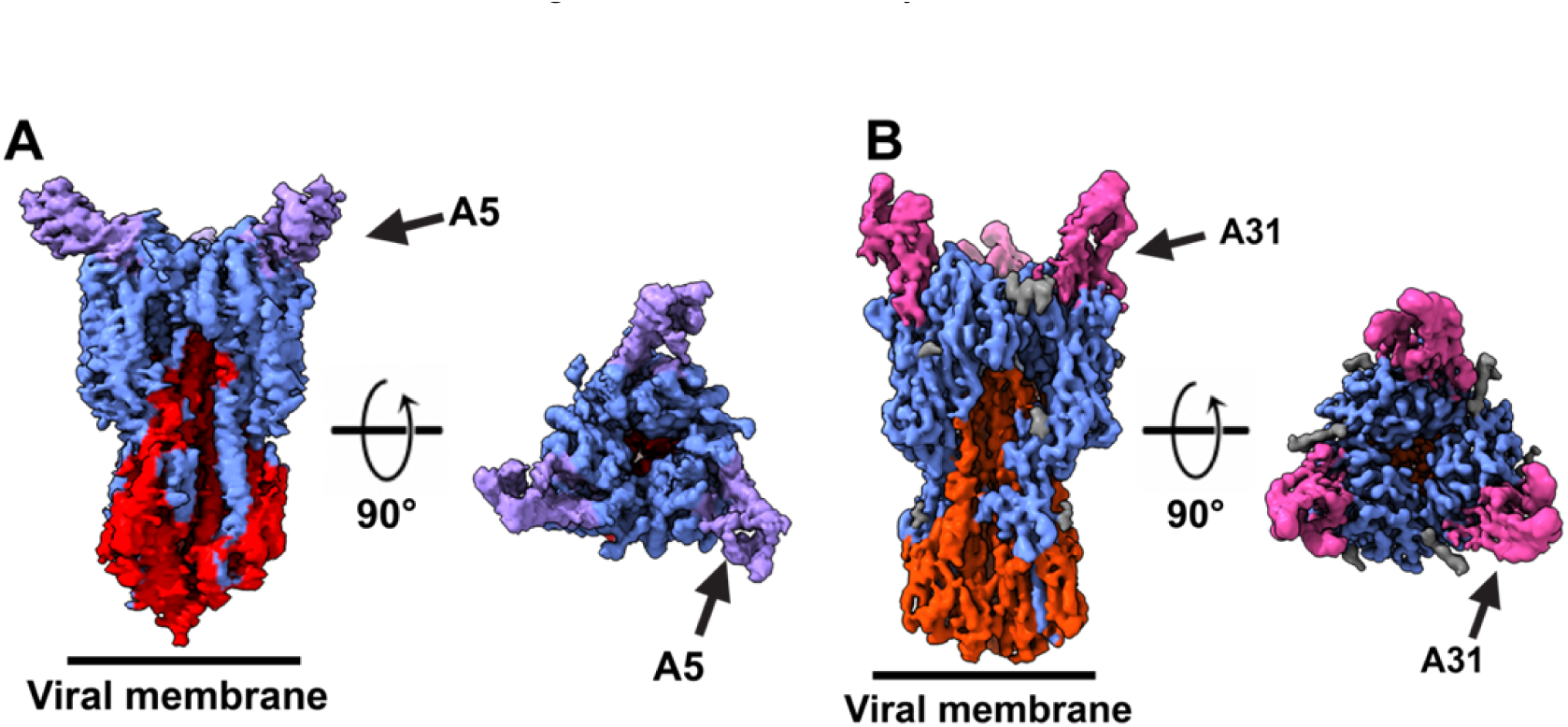
Affimer-HA complex determination by cryo-EM. Cryo-EM maps of the HK68 H3-A5 **(A)** and HK68 H3-A31 **(B).** The HK68 H3-A5 map was filtered using deepEMhancer to enable fitting of an HA trimer (PDB: 4FNK) and an Alphafold2 generated model of A5 (28). HA1 is blue, HA2 is red, A5 is purple, A31 is pink, and glycans are grey.

### The structural and functional basis of HA-A31 interactions

Our two cryo-EM structures showed that both Affimer molecules (A5 and A31) specifically interact with and/or obstruct the RBD of H3 (Fig. 3A and B). The resolution of the H3-A31 average enabled model building and mapping of interactions between A31 and HA. In this model, the two variable peptide regions of A31 appear to straddle the 130-loop of HA (Fig. 4A), which is directly involved in sialic acid building (Fig. 4B) (29). This suggests that the mechanism of action of A31 (and similarly, of A5), is through competition with sialic acid binding. The interactions of A31 with HA highlight multiple contact points directly (G135, W153, K156, W222 and S227), or in immediate proximity (R141, G146, R220 and R255) to HA sialic acid interacting residues (Figs. 4A and B) (29).

**Figure 4.**
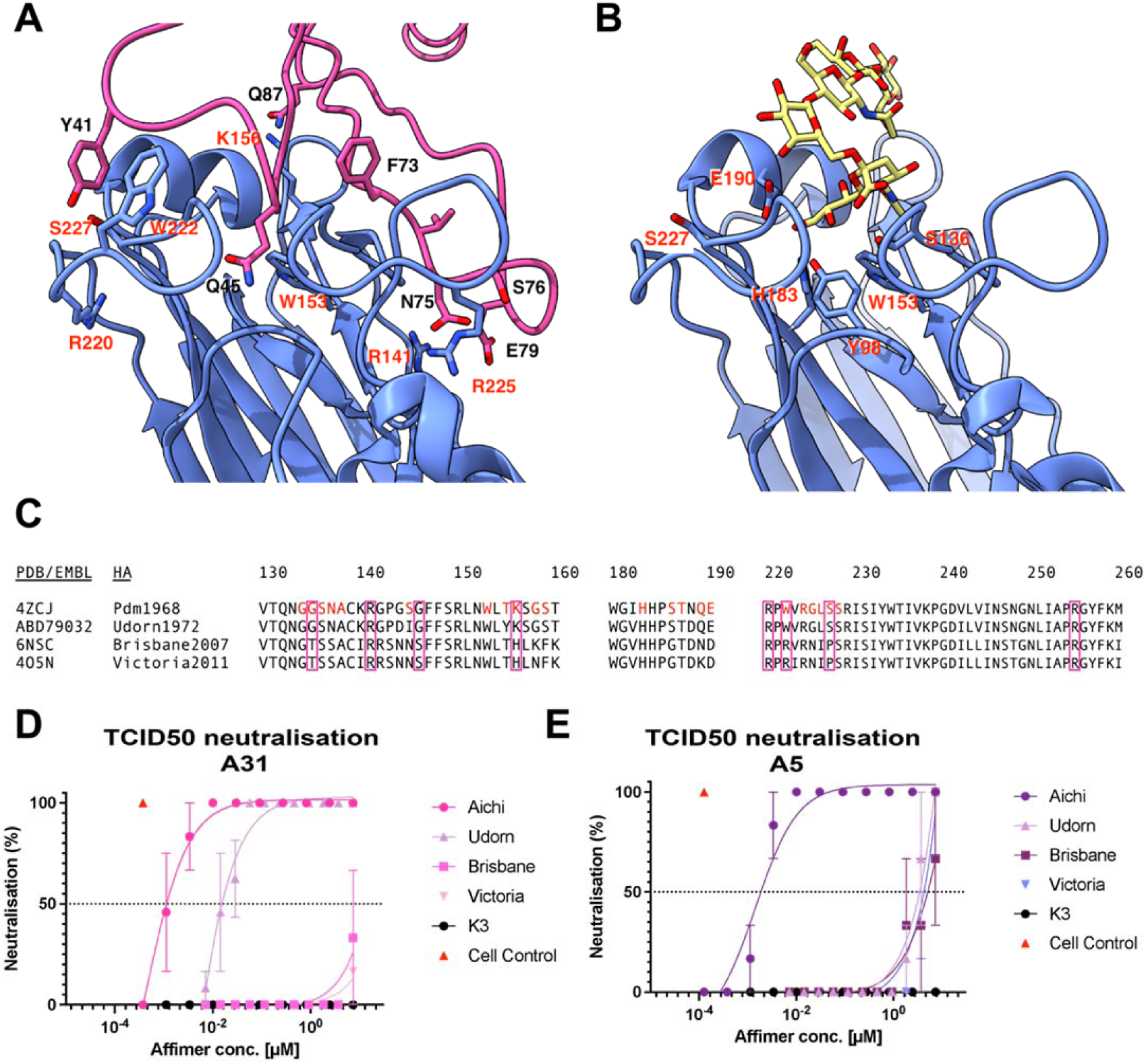
Affimers A5 and A31 target the RBD of A/Aichi/1968 HA and can neutralise related strains. **(A)** Interacting residues of A31 variable loops with the RBD of HK68 HA. **(B)** Sialic acid (yellow) docked into the RBD of HA1 with interacting residues labelled. **(C)** Multiple sequence alignment of the RBD of a range of H3N2 viruses. Directly interacting residues with sialic acid for A/Aichi/1968 (H3N2) HA are highlighted in red. Purple boxes show interacting residues with A31. **(D-E)** 100 x TCID_50_ assay in which neutralisation of either A31 **(D)** or A5 **(E)** was tested against a range of IAV strains in a concentration-dependent manner (7.41 μM-125.5 pM). In D-E, data are mean and error bars represent standard deviation (from two biological repeats, each performed in triplicate).

To confirm the H3-A31 interactions, a panel of H3N2 strains were selected with similar, or distinct residues in the region of H3-A31 interactions. The selected strains included the closely related virus *A/Udorn/1972 (H3N2)*, which exhibits limited adaptions in the sialic acid interacting site (with the exception of G144D, S145I and T155Y, all other residues within this site are identical to A/Aichi/1968 (H3N2); Fig. 4C). Additionally, the more recent circulating strains *A/Brisbane/2007 (H3N2)* and *A/Victoria/2011 (H3N2)* were selected to provide distinct residues in the interacting region.

Affimer molecules were assessed against these strains using TCID_50_ assays (Fig. 4D and E). A31 maintained high potency, although somewhat decreased, when challenged against *A/Udorn/1972 (H3N2)* (TCID_50_ of 18.22 nM vs. 0.44 nM against *A/Aichi/1968 (H3N2)*; Fig. 4D). This could be a consequence of the slight change in the binding pocket due to G144D, S145I and T155Y mutations (Fig. 4C and Sup. Fig. 3B). On the other hand, a decrease in potency was observed when using A5 to neutralise *A/Udorn/1972 (H3N2)* (TCID_50_ of 3 µM vs. 1.96 nM against *A/Aichi/1968 (H3N2);* Fig. 4E and Sup. Fig. 3B). This suggests that one or more of the Udorn-specific mutations when compared to Aichi (G144D, S145I and/or T155Y) are important for A5 binding to HA. When *A/Brisbane/2007 (H3N2)* and *A/Victoria/2011 (H3N2)* were challenged with A5 and A31, TCID_50_ values were ≥ 3 µM, indicating A5 and A31 inhibit these strains, but the potency is three orders of magnitude lower compared to *A/Aichi/1968 (H3N2*) (Figs. 4D and E, and Sup. Figs. 3C and D). Overall, these results validate the cryo-EM observations: as expected from RBD-interacting molecules, both Affimers target a highly variable epitope, and as such they lose potency when challenged against differing HAs. However, A31 does present some degree of protection against IAV strains closely related to *A/Aichi/1968 (H3N2)* in terms of their RBD.

### Affimers prevent viral entry through interaction with the receptor binding site

HA is the main target of neutralising antibodies due to its higher abundance compared to NA. Given the different functions of HA during the IAV replication cycle, neutralising antibodies appear to act by either blocking receptor binding, preventing key conformational changes required for successful fusion of viral and cellular membrane, or by inhibiting the release of progeny virions (30). Of note, the most potent antibodies typically inhibit receptor binding by blocking the RBD on HA1, while broadly neutralising antibodies typically target the fusion domain in HA2 (31), although some broadly neutralising antibodies to the receptor binding site of NA have been recently isolated (32). To further validate RBD binding and to confirm the mechanism of inhibition of A5 and A31, a series of assays were conducted (Sup. Fig. 8).

First, to confirm that A5 and A31 inhibit sialic acid binding, a classical IAV hemagglutination inhibition assay was performed (Sup. Fig. 8A). In this assay, human red blood cells (RBCs) are incubated with a series of candidates for inhibiting hemagglutination. In the absence of hemagglutination, RBCs accumulate at the bottom of the well, resulting in a red-coloured dot. When IAV alone is added to RBCs, hemagglutination occurs, inducing clumping of RBCs which can no longer accumulate at the bottom of the well, resulting in the absence of a dot. When IAV-induced hemagglutination is blocked, for example due to the presence of an Affimer blocking the interaction with sialic acid, RBCs settle to the bottom of the well resulting in a red dot. When the hemagglutination of IAV A/Aichi/1968 was tested in the presence of A5 and A31, no hemagglutination was detected, even at the lowest concentration tested (223 nM or 3.125 µg/mL; Fig 5A), confirming previous findings that both Affimer molecules A5 and A31 prevent IAV binding to host cell sialic acids.

**Figure 5.**
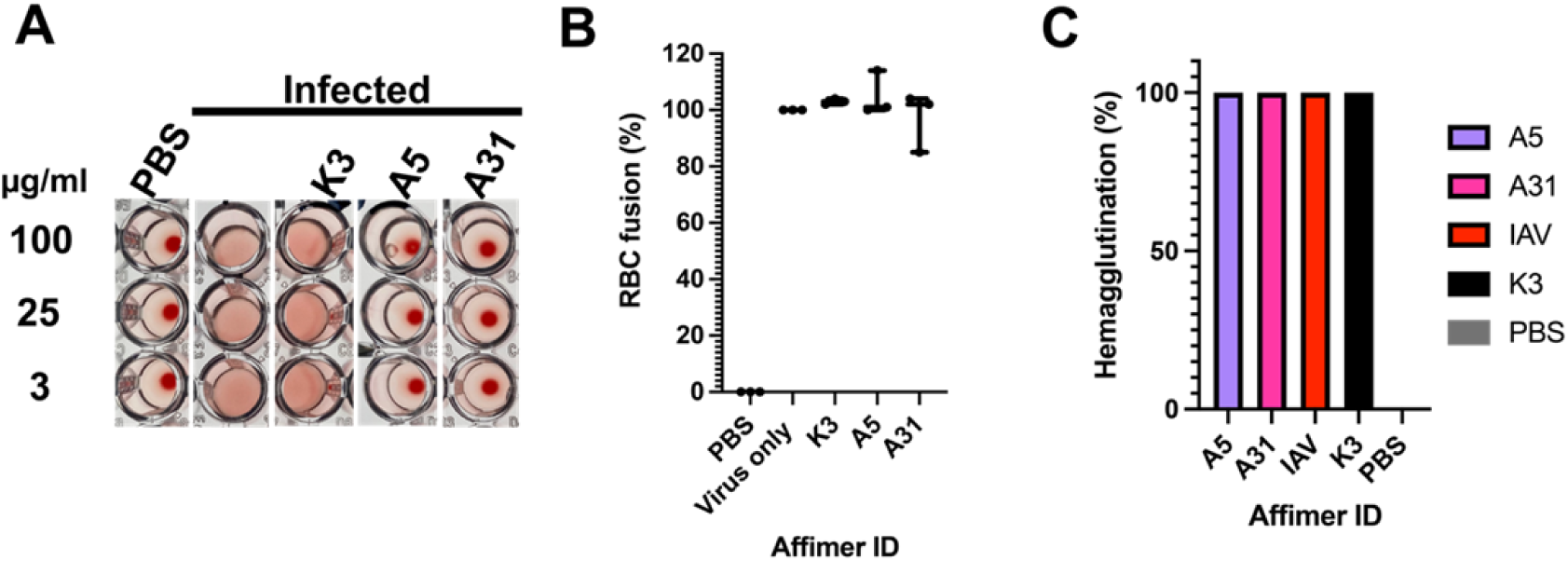
Confirmation of the mechanism of inhibition of A5 and A31. **(A)** Classical hemagglutination assay in which influenza virus A/Aichi/1968 (H3N2) (4 HA units), was challenged with either: A5, A31; or negative controls A12, K3 (100 µg/ml, 25 µg/ml or 3 µg/ml), before adding 1% v/v hRBC. hRBC incubated with PBS in the absence of virus was used as a positive control for hemagglutination. **(B)** An adapted fusion assay (22), in which hRBCs were challenged with virus or virus-Affimer complexes and exposed to low pH (pH 4) to determine if fusion occurs (release of NAPDH), measured at A340 nm. % fusion was determined by analysing against virus only control. Data are mean and error bars represent standard deviation (from two biological repeats, each performed in triplicate). **(C)** An egress assay in which virus infection was synced on ice before allowing a single round of infection to occur (4 hrs). Affimers were then added and infection allowed to continue for a further 18 hrs before supernatant was used to assess hemagglutination of hRBCs. Two biological repeats were carried out for each assay.

We then tested whether A5 and A31 presented additional mechanisms of inhibition, such as blocking fusion or interrupting viral egress. To verify if Affimer molecules inhibited HA fusion, an *in vitro* assay was adapted in which RBCs were incubated on ice with/without IAV to synchronise binding (22) (Sup. Fig 8B). Control or Affimer molecules were then added to RBCs incubated with IAV, before exposing the virus to a fusion buffer (pH 5.0) for 30 mins. In the absence of a fusion inhibitor, structural re-arrangements of HA to a post-fusion state induce RBC lysis, causing the release of NADPH, which can be quantified indirectly by an increase in the absorbance at 340 nm. The presence of a fusion inhibitor prevents NADPH from being released to the media and the absorbance at 340 nm does not increase. In the presence of A5 and A31, RBC lysis was also observed, comparable to levels of the non-specific Affimer K3, indicating the Affimer molecules did not inhibit fusion (Fig. 5B).

Finally, we explored if A5 and A31 affected viral egress. In this egress assay, cells were pre-incubated with virus for 4 hrs, prior to the addition of A5, A31 or control, enabling entry and a single round of infection to occur in the absence of Affimers or control. Following addition of Affimer molecules or control, infection was allowed to continue for a further 18-20 hrs, before a hemagglutination assay was carried out. If egress was inhibited, no virions would be present in the media and therefore hemagglutination would be observed and RBCs would settle to the bottom of the well (Sup. Fig. 8C). Our results showed neither A5 nor A31 inhibited hemagglutination, therefore they had no effect on viral egress (Fig. 5C). Overall, these results suggest that the mechanism of A5 and A31 inhibition is based solely on their blocking of sialic acid-HA binding by interacting at or around the HA RBD.

## Discussion

To better prepare for future pandemics, multiple strategies are required. Alongside preventive vaccination approaches, mAbs have been favoured for therapeutic intervention, due to high potency viral neutralization and promising animal and clinical efficacy (30). However, due to the nature of pandemic outbreaks, global requirements for rapid manufacturing alongside the cost of production and distribution is a major bottleneck for both vaccines and antibodies (33,34). Furthermore, resistance rapidly makes therapeutics such as mAbs and antivirals ineffective (3). This highlights the requirement for a variety of interventions, particularly those which can be rapidly and cheaply produced under the threat of emerging pandemic variants. Several alternative therapeutics are under development (35), with some showing success in a clinical setting, such as the DARPin-based Ensovibep, which is in phase II clinical trials directed against SARS-CoV-2 (36). These alternatives, including for example nanobodies (37), will likely complement current treatment and perhaps overcome some of the current limitations, such as the disadvantages associated with mAb production and treatment (38).

Here we showcase the isolation and characterization of Affimer molecules, an alternative to mAbs rapidly being shown to be an effective diagnostic and therapeutic against not only oncological, but more recently viral targets (13–15,39), as tools against IAV. We carried out a small-scale screen from a library of ∼10^10^ Affimer molecules using only the monomeric form of HA (A/Aichi/1968), identifying 34 unique Affimer molecules with different functionalities and binding specificities.

The two monovalent Affimer molecules that were characterised in detail and highlighted here target the RBD, impeding interaction with sialic acid receptors that is necessary for infection of the host cell. We demonstrate that the Affimers not only make high affinity interactions with HA (K_d_ <4 nM), but also show high protection against IAV *in vitro* (TCID_50_<2 nM). The affinity of these Affimer molecules for the target protein alongside their potency of virus neutralization is similar to the profile of potent mAbs raised against influenza viruses (18–24). Since Affimer molecules have already shown good promise as therapeutics, our results pave the way for the use of Affimers as anti-viral therapeutics against IAV in a clinical setting.

However, Affimers A5 and A31 present low potency against more recent IAV strains compared to the one that they were screened against. This suggests that Affimers A5 and A31 may not be as effective *in vivo* against these strains. One plausible explanation for this reduced inhibition lies in the presence of additional N-glycosylation sites, N133 and N144, which were acquired and subsequently maintained through evolutionary processes in all H3N2 strains from 1974 onwards (40). Glycosylation at these sites could potentially interfere with the binding affinity and overall effectiveness of A5 and A31 against the A/Brisbane/2007 and A/Victoria/2011 strains of H3N2 influenza since they are situated near the regions where both Affimers interact with HA. Of note, the breadth of some mAbs is based on inserting a single complementarity-determining region into the conserved receptor binding site, minimising contact with the surrounding more hypervariable sites (41,42). A potential way to isolate Affimers targeting only the conserved receptor binding site would be to perform phage display panning rounds employing HA molecules from different strains, allowing isolation of broad-spectrum Affimers.

Importantly, Affimer molecules have the capacity to overcome many disadvantages that are encountered with therapeutic mAbs. Their small size increases their solubility and rapid tissue penetration, which can often be a setback for mAbs in a clinical setting (38). Affimers typically display a high degree of temperature stability, which might also enable alternative routes of administration, such as inhalation, as has been described for other highly stable protein scaffolds (43,44). Furthermore, Affimer molecules lack an Fc-region, likely preventing antibody dependent enhancement (ADE) effects, a potential side-effect of mAbs in patients with inflamed lungs (45), a typical symptom of aggressive respiratory infections (45).

Furthermore, the multimerization of small antibody-like proteins can further enhance individual molecules effectivity (36,46), providing the opportunity to increase avidity for target viral proteins, in the case of same-target multimeric molecules. This approach also provides the potential to improve potency through multimeric molecules targeting multiple immunogenic sites of a viral protein at once, such as the RBD and conserved stem of HA, raising the fitness barrier for escape mutants. To improve half-live, bi-specific Affimer reagents that bind to human serum albumin can also be created, as clearance is an issue for small biologics. We anticipate that the presented workflow for Affimer development could be applied to any future circulatory or more crucially emerging pandemic strain of IAV, and also broadly to other diseases of concern.

Additionally, the Affimers described here could be used as diagnostic tools to detect IAV. Affimers have already been employed for ELISAs to diagnose plant virus diseases (14) and for colorimetric diagnostic tests against Crimean-Congo hemorrhagic fever virus (13). Therefore, broad-spectrum Affimers as described above could be used within these systems to detect viral infection. Furthermore, an Affimer-Enzyme-Inhibitor Switch Sensor has been developed (47) that would allow multiplexed detection of respiratory viruses when IAV-specific Affimers are combined with Affimers against other respiratory viruses (e.g. SARS-CoV-2, human respiratory syncytial virus or influenza virus B).

Overall, we have shown that high-affinity and potently neutralizing Affimer molecules can be isolated against influenza A virus, and we have established a workflow to characterise these molecules that could be performed in a matter of weeks, without the requirement of whole antigen, immunization of animals or access to patient serum. Ultimately, fast-track development strategies of stable and potent inhibitors are critical to raise the global preparedness level towards novel pandemic viruses.

## Methods

### IAV proteins

The monomeric HA protein employed to select Affimer molecules was a His-tagged monomeric IAV HA derived from A/Aichi/2/1968 (H3N2; SinoBiological Cat: 11707-V08H). To assess affinity through ELISA and SPR, both monomeric HA and HA1 (Cat number 11707-V08H1) from A/Aichi/2/1968 were used (both were His-tagged). For cryo-EM structure determination, a trimeric HA from A/HongKong/1968 was generated and purified as previously described (48).

### Cells and viruses

H3N2 IAVs were propagated in MDCK cells including A/Aichi/2/1968 (ATCC® VR-1680), A/Udorn/307/1972 (provided by Dr. Lee Sherry, University of Leeds), A/Brisbane/10/2007 (BEI resources, NIAID, NIH: NR-12283), and A/Victoria/361/2011 (BEI resources, NIAID, NIH: NR-44022). Cells were cultured in Minimum Essential Media (MEM, Sigma) supplemented with 1% FBS and 1x antibiotic antimycotic, which was also supplemented with 10% DMSO.

### Selection of HA-specific Affimer molecules by phage display

To select Affimer molecules through phage display, phage libraries were utilized and subjected to three rounds of panning against the monomeric HA protein of A/Aichi/2/1968 (H3N2), as previously described (12). The target protein was immobilized on streptavidin coated plates and pre-panned phage were incubated overnight before washing with increasing stringency. Isolated phages were then amplified for phage ELISA.

### Screening of isolated Affimer molecules

Phage ELISA screening was performed, as previously described (12), on randomly selected clones from the final panning round of Affimer selection. Screening enabled the positive selection for final evaluation and characterisation. To enable a small-scale assessment of IAV directed Affimer-based therapeutics, an arbitrary cut-off point of >0.5 absorbance reading (at 620 nm) was implemented. Successful candidates were Sanger sequenced and unique Affimers expressed, purified, and characterised.

### Affimer production and characterisation

After sequencing, the ORFs of unique Affimers were subcloned into a pET11a expression vector. The resultant Affimers were tagged with an N-terminal 8x His-tag and cysteine for functionalization via bacterial expression (*E. coli* strain Rosetta 2). The Affimers were purified using Ni^2+^-NTA affinity chromatography following previously described methods (12).

### Cell viability assay

To determine cell viability, an ATPlite^TM^ assay was performed following the Perkin Elmer 1- step protocol with slight modifications. Initially, MDCK or A549 cells were seeded in 96-well clear-bottom tissue culture plates at densities of 3 x 10^4^ cells/well and 1.25 x 10^4^ cells/well, respectively, and incubated at 37 °C with 5% CO2 for 24 hours. Once the cells reached 80- 90% confluence, the growth medium was removed, and the cells were washed twice with PBS. Next, Affimers at 100 µM (the highest concentration employed for any assay) were added to triplicate wells. All drug conditions were prepared in infection media. The cells were then incubated at 37 °C with 5% CO2 for 48 hrs. To measure luminescence, 50 μL of mammalian cell lysis solution was added and incubated on a plate shaker for 5 minutes, followed by the addition of 50 μL of substrate solution and incubation for a further 5 minutes. The plate was then dark-adapted, and luminescence was measured at 510 nm using an ELISA plate reader.

### Virus neutralization assay

The capacity of Affimer molecules to neutralize viruses was determined by TCID_50_ assay, by crystal violet staining of protected cells in the presence of IAV. Briefly, Affimer candidates were serially diluted from 7.41 μM-125.5 pM in cell culture media (1%-FBS-MEM) (in duplicates with 3 independent repeats). The diluted Affimer candidates were then exposed to various IAV strains at 100 x TCID_50_ in 1%-FBS-MEM. Affimer/virus mixtures were transferred onto 80% confluent MDCK cells. Controls included MDCK cells exposed to Affimer molecules only, cells exposed to virus incubated with a non-IAV specific Affimer (K3; as a negative control), cells exposed to virus only (to determine maximal cytopathic effect), and cells incubated with medium only (to determine the baseline state of cells). The plates were incubated for 2-5 days (strain-dependent) at 37°C, and the cytopathic effect was determined by staining with crystal violet solution (0.5% crystal violet diluted in 37% formaldehyde solution and PBS; all reagents from Sigma Aldrich) for 10 minutes, followed by washing plates with PBS. Wells were observed for complete protection indicated by an intact blue/violet cell layer, or partial protection in case of ∼50% intact cell layer.

### Surface plasmon resonance affinity determination of Affimer molecules

Using a Biacore 3000 (GE Healthcare), biotinylated Affimers were immobilized on a Sensor Chip SA (GE Healthcare) through streptavidin-biotin interaction. Affimers were diluted to a concentration of 100 nM in PBS and injected into the respective flow cells at a flow rate of 5 μL/min until the surface density reached 100 response units. A flow cell was left unoccupied as a reference surface. Monomeric HA was diluted in PBS to various concentrations and injected at a flow rate of 20 μL/min for 120 seconds. BIAevaluation software was employed for double-referencing analysis. Affinity and kinetic constants were determined using a Langmuir 1:1 binding model and steady-state affinity models.

### ELISA based affinity determination of Affimer molecules

Either the monomeric HA or HA1 head domain (both from Aichi and 5 μg/mL per well) were immobilized on Maxisorb plates (Nunc) overnight at 4 °C, followed by blocking with 1x casein blocking buffer (Sigma) for 4 hours at room temperature. After washing once with PBS, the plates were incubated with a range of concentrations (7.41 μM-125.5 pM) of biotinylated anti-HA Affimer for 1 hour at room temperature. Subsequently, the plate was washed with PBST and the bound anti-HA Affimer molecules were detected with a 1:1000 dilution of HRP-conjugated streptavidin (Pierce) for 1 hour at room temperature. After washing the plates 10 times with PBST, Affimer molecule binding was visualized with TMB (Seramun) and measured at 405 nm.

### Mechanism of inhibition determination by Affimer molecules

The Affimer molecules were serially diluted in PBS in 96-well U-bottom plates at different concentrations (7.41 μM-125.5 pM (in duplicates, 3 independent repeats)) and exposed to pre-determined-4 HA units of H3N2 (A/Aichi/2/1968) diluted in PBS. After incubating the mixture for 45 minutes at 37°C, 2% v/v hRBC (Cambridge Bioscience, RBC1DC4CIT03- XSXX) were added at 1:1 vol/vol ratio and incubated at room temperature for 1 hour. The hemagglutination effect was then observed visually.

The RBC fusion assay was adapted from previously described methods (22), which involved incubating 1% v/v hRBC and H3N2 virus (A/Aichi/2/1968) in a 1:1 ratio on ice for 30 mins, followed by adding different concentrations of Affimer molecules (7.41 µM, 1.48 µM, 148.15 nM) including a control Affimer (K3). Samples were spun down at 4,000 *x g* for 3 mins, supernatant aspirated and 200 μL of buffered solution (15 mM citric acid (pH 5.0), 150 mM NaCl2) before incubating at 37 °C for 30 mins. Samples were spun at 4,000 *x g* and supernatant harvested and the lysis of RBCs was measured by the presence of NADPH (Absorbance at 340 nm).

For the MDCK cell assay, the cells were first infected with an MOI of 5, H3N2 (A/Aichi/2/1968). Affimer molecules or control Affimer K3 were added at 4 hours post-infection. At 8 hours post-infection, the supernatants were collected, and nascent virus was assessed using a hemagglutination assay.

### Cryo-electron microscopy sample preparation of HA

In cryo-EM grid preparation, HA shows strong preferred orientation which can complicate cryo-EM processing (26). We first tested whether our custom-built setup for fast grid preparation would reduce preferred orientation of HA, to enable structure determination of HA in the absence of Affimers and HA-Affimer complexes. A schematic of the setup used for this experiment is given in Supplementary Fig. 4A. The HA sample was used at 2.9 mg/ml (17.5 µM). Quantifoil 300 mesh copper R1.2/1.3 grids were used after glow discharge in a Cressington 208 carbon coater with glow-discharge unit for 99 s at 0.1 mbar air pressure and 15 mA.

The sample was injected into a gas-dynamic virtual nozzle made of PDMS (49), at a liquid flowrate of 5.2 µL/s. A spray of the sample was generated by applying an N_2_ gas pressure of 2 bar to the nozzle’s gas inlet. The spray was allowed to stabilise for 0.8 s, then the grid was moved through the spray at 1.9 m/s. The distance between spray nozzle and grid was 10 mm at the point of sample application, and the distance between sample application and freezing was 22 mm. This resulted in a residence time of ∼ 12 ms for the sample on the grid. The settings for grid preparation of all samples are summarized in Supplementary Table 1.

### Cryo-electron microscopy of sample preparation HA-A5

For the HA-A5 complex, we modified the setup to allow for rapid mixing and freezing. Previous experiments using pre-mixed HA and Affimer A5 showed aggregated particles that were not amenable to structure determination. Therefore, we chose in-flow mixing with a time delay of 700 ms between mixing of HA and A5 and freezing. The method has been described in detail elsewhere (50), and a schematic of the setup is shown in Supplementary Fig. 4C. Self-wicking grids supplied by SPT Labtech were used after glow discharge in a Cressington 208 Carbon coater with glow-discharge unit for 80 s at 0.1 mbar air pressure and 15 mA.

The HA sample (5.9 mg/mL, 35 µM) and the A5 sample (1.5 x molar excess Affimer A5 and 0.2 % octyl glucoside) were loaded into the grid preparation system independently. Mixing of HA and A5 was initiated in a mixing unit upstream of the spray nozzle. The binding reaction took place between the mixing unit and the spray nozzle, in the “delay-line”, which was a 20 mm segment of tubing with 381 µm inner diameter. The samples travelled through the system with a combined flowrate of 4.2 µL/s. Assuming laminar flow, this led to a median reaction time of 700 ms. The reaction time for 87 % of all particles was between 580 ms and 1000 ms. The flow through the setup was initiated for 1 s before sample application and plunge-freezing.

### Cryo-electron microscopy sample preparation of HA-A31

For the HA-A31 complex, 2.9 mg/ml HA (17.5 µM) was pre mixed for 15 minutes at room temperature with 1.5 x molar excess of Affimer A31 and 0.1% octyl glucoside, before vitrifying by rapid cryo-EM grid preparation. A schematic of the setup is given in Supplementary Fig. 4E. A. Self-wicking grids (SPT Labtech) were used after glow discharge in a Cressington 208 carbon coater with glow-discharge unit for 80 s at 0.1 mbar air pressure and 15 mA.

The sample was injected into a gas-dynamic virtual nozzle made of PDMS (49) at a liquid flowrate of 4.2 µL/s. A spray of the sample was generated by applying an N_2_ gas pressure of 2 bar to the nozzle’s gas inlet. The spray was allowed to stabilise for 0.8 s, then the grid was moved through the spray at 1.5 m/s. The distance between spray nozzle and grid was 12 mm at the point of sample application, and the distance between sample application and freezing was 22 mm. This resulted in a residence time of ∼ 15 ms for the sample on the grid.

### Cryo-electron microscopy data collection

For HA in the absence of Affimers, movies were collected using a Titan Krios cryo-TEM (Thermo Fisher Scientific) operating at 300 keV and equipped with a K2 Direct Electron Detector (Gatan). Data were acquired using the EPU 2 software (Thermo Fisher Scientific). Movies were collected in electron counting mode at 130,000x corresponding to a calibrated pixel size of 1.07 Å/pix over a defocus range of -3 to -5 μm.

For HA-A5 and HA-A31, movies were collected using a Titan Krios Cryo-TEM (Thermo Fisher Scientific) operating at 300 keV and equipped with a Falcon 4 Direct Electron Detector (Thermo Fisher Scientific). Cryo-EM data were acquired using the EPU 2 software (Thermo Fisher Scientific). Movies were collected in electron counting mode, over a defocus range of -2 to -4 μm and at a nominal magnification of 96,000x which corresponded to a calibrated pixel size of 0.83 Å/pix.

### Cryo-electron microscopy data processing of HA

For the dataset of HA in the absence of Affimers, all processing was done in RELION 3.1 (51). Movies were imported and underwent beam-induced motion correction using MotionCor2 (52). Then, the contrast transfer function (CTF) of each micrograph was estimated using gctf (53), crYOLO 1.6.1 (Sphire) was used to automatically pick particles on the motion-corrected micrographs using the weights from its general model and a picking threshold of 0.1. 230,312 particles were picked and extracted into a 280-pixel box re-scaled to 140 pixels. After 2D classification, 151,911 “good” particles were selected. A subset of 50,000 particles was chosen to generate an initial model (with C3 symmetry). This initial model was used for 3D classification of the full set of selected particles with 6 classes over 70 iterations and with C1 symmetry. Particles contributing to the best class (129,501 particles) were reextracted in a 300-pixel box and underwent 3D refinement with C3 symmetry. Two rounds of Bayesian polishing and CTF refinement produced the final structure with a resolution of 4.3 Å.

### Cryo-electron microscopy data processing of HA-Affimer complexes

Image processing for both datasets began with the same steps. Movies were imported into RELION 3.1 (51) and underwent beam-induced motion correction using RELION’s own implementation (52). Then, the contrast transfer function (CTF) of each micrograph was estimated and corrected for using CTFFIND-4.1 (54). crYOLO 1.6.1 (Sphire) was used to automatically pick particles on the motion corrected micrographs using the weights from its general model and a picking threshold of 0.1. 200,759 particles were picked on the H3-A31 dataset and 275, 510 particles were picked on the H3-A5 dataset. Particles for both datasets were extracted into a 280-pixel box re-scaled to 100 pixels.

All H3-A5 processing was performed using Relion 3.1. Following extraction, particles underwent one round of 2D classification and the 227,015 particles in the HA-containing classes were taken forward for 3D classification. Only one 3D class contained density for HA with bound Affimers, so particles in this class were extracted in a 280-pixel box re-scaled to 200 pixels. These particles were refined with C3 symmetry to a map with 5.6 Å. Following two rounds of bayesian polishing and CTF refinement, the final map with C3 symmetry had a final map of 4.4 Å according to the gold standard half-map criteria at a 0.143 cut-off.

From here, the datasets were treated differently. H3-A31 particles were imported into cryoSPARC (55) and underwent one round of 2D classification. Classes containing HA were taken forward for *ab initio* 3D model generation and particles classified into models containing HA were taken back into RELION and re-extracted in a 400-pixel box without re- scaling. The unbinned particles were then processed by cryoSPARC using the algorithm for 3D non-uniform refinement without applied symmetry and then with C3 symmetry. Particles were imported into RELION for Bayesian polishing, then back to cryoSPARC for a second round of non-uniform refinement and global sharpening. This resulted in final map with a global resolution of 3.41 Å according to the gold standard half-map criteria at a 0.143 cut-off.

Overviews of the image processing workflows and statistics are shown in Sup. Figs. 5-7, and Sup. Table 2.

### Molecular modeling of H3-A31

AlphaFold models of Affimer molecules A5 and A31 were first generated (28). HK68 (PDB: 4FNK) HA and the Alphafold Affimer models were rigid-body fitted into the cryo-EM maps produced using the UCSF Chimera ‘fit in map’ tool (56). H3-A31 was then modelled by first improving the fit in Coot (57), before utilizing Namdinator (58). To aid model fitting around the Affimer region, the final H3-A31 map was sharpened using DeepEMhancer (59) as implemented in COSMIC2 (60). This map was used alongside cryoSPARC maps to improve confidence in A31 modelling. The final model was refined in Phenix and iteratively improved in Coot until the model was considered satisfactory. Figures were generated using UCSF Chimera and UCSF ChimeraX (61).

## Data Availability

The cryo-EM maps for H3 HA, H3-A5 and H3-A31 complexes have been deposited in EMDB (accession numbers EMD-18137, EMD-17725 and EMD-17724). The atomic model for the H3-A31 complex has been deposited in PDB (accession number: 8PK3).

## Acknowledgements

We thank the Astbury Biostructure Laboratory electron microscopy facility, the Protein Interactions facility (Iain Manfield) for support in SPR experiments, Amy Turner from the University of Leeds for support purifying Affimers, and Xueyong Zhu and Chung-Chun D. Lee at The Scripps Research Institute, for providing the trimeric HA protein used in all cryo-EM experiments. Influenza virus strains A/Brisbane/10/2007 and A/Victoria/361/2011 were obtained through BEI Resources, NIAID, NIH (reference numbers NR-12283 and NR- 44022).

